# Deep learning predicts DNA methylation regulatory variants in specific brain cell types and enhances fine mapping for brain disorders

**DOI:** 10.1101/2024.01.18.576319

**Authors:** Jiyun Zhou, Daniel R. Weinberger, Shizhong Han

## Abstract

DNA methylation (DNAm) is essential for brain development and function and potentially mediates the effects of genetic risk variants underlying brain disorders. We present INTERACT, a transformer-based deep learning model to predict regulatory variants impacting DNAm levels in specific brain cell types, leveraging existing single-nucleus DNAm data from the human brain. We show that INTERACT accurately predicts cell type-specific DNAm profiles, achieving an average area under the Receiver Operating Characteristic curve of 0.98 across cell types. Furthermore, INTERACT predicts cell type-specific DNAm regulatory variants, which reflect cellular context and enrich the heritability of brain-related traits in relevant cell types. Importantly, we demonstrate that incorporating predicted variant effects and DNAm levels of CpG sites enhances the fine mapping for three brain disorders—schizophrenia, depression, and Alzheimer’s disease—and facilitates mapping causal genes to particular cell types. Our study highlights the power of deep learning in identifying cell type-specific regulatory variants, which will enhance our understanding of the genetics of complex traits.

**Teaser:** Deep learning reveals genetic variations impacting brain cell type-specific DNA methylation and illuminates genetic bases of brain disorders

## Introduction

DNA methylation (DNAm) is essential for brain development and function, and its aberrations are implicated in neurological and psychiatric disorders(*1–4*). Genetic association studies have identified genetic variations associated with DNAm levels in the human brain, known as DNAm quantitative trait loci (mQTLs)(*5–9*), which may illuminate causal genetic variations within risk loci identified by genomewide association studies (GWAS)(*10, 11*). However, the majority of those studies were conducted with bulk tissues and may not capture cell type-specific mQTLs, thereby limiting their utility to uncover risk variants that specifically act in disease-relevant cell types(*12–14*). Advances in single-cell technologies have enabled profiling of DNAm at single-nucleus resolution in the human brain(*15–17*), providing an opportunity to identify cell type-specific mQTLs through genetic association studies. However, the current cost of these technologies is still too expensive to generate a sufficient sample size for robust statistical power. Additionally, genetic association studies face challenges in identifying functional variants that drive DNAm levels due to extensive linkage disequilibrium (LD) across the genome.

As a complementary approach to population-based QTL studies, deep learning techniques have emerged as a promising tool for predicting the effects of genetic variations on various molecular traits(*18*), such as gene expression(*^19, 20^*), chromatin marks(*21, 22*) and DNAm levels(*23*). These techniques first build prediction models for molecular traits based on local DNA sequences and then estimate the impact of genetic variations on these traits by comparing the predicted levels of molecular traits between the two DNA sequences of different alleles. Deep learning-based approaches offer advantages over traditional QTL studies as they do not rely on population-based samples and are not confounded by LD(*23*). Instead, these approaches predict regulatory variants by assessing their impact on DNA motifs associated with molecular traits. In a previous study, we developed a deep learning model INTERACT, which integrates convolutional neural network (CNN) with the attention mechanism of transformer to predict variant effects on DNAm levels in bulk brain tissues(*23*). Our study demonstrated the superiority of INTERACT over a standard CNN model, but the limitation was that our previous model was trained on bulk brain samples, limiting its ability to detect cell type-specific effects.

Here, we extend INTERACT to predict DNAm regulatory variants in specific brain cell types utilizing existing single-nucleus DNAm data from the human brain. We show that INTERACT models, trained for each cell type, accurately predict cell type-specific DNAm profiles and uncover DNA motifs and transcription factors that may underline these profiles. Furthermore, INTERACT predicts cell type-specific DNAm regulatory variants, which reflect cellular context and enrich heritability of brain-related traits in relevant cell types. Importantly, we demonstrate that incorporating predicted variant effects and DNAm levels of CpG sites enhances the fine mapping of risk loci for three brain disorders---schizophrenia, depression and Alzheimer’s disease---and facilitates mapping potential causal genes to particular cell types. Our study highlights the power of deep learning in identifying regulatory variants in specific cell types, which will enhance our understanding of the genetic underpinnings of complex traits.

## Results

### INTERACT model for predicting cell type-specific DNAm levels

We designed INTERACT, a deep learning model that combines CNN and the attention mechanism of transformer, to predict DNAm levels of CpG sites from local DNA sequences (**Figure 1A**). To train cell type-specific INTERACT models, we utilized an existing single-nucleus DNAm dataset containing 4,137 nuclei from the human prefrontal cortex(*16*), which were clustered into 13 cell types, including four excitatory neuron subtypes (L2/3, L4, L5, and L6), four inhibitory neuron subtypes (Ndnf, Vip, Pvalb, and Sst), and five non-neuronal subtypes (astrocyte, oligodendrocyte (ODC), oligodendrocyte progenitor cell (OPC), microglia, and endothelial cell). Given the model’s complexity (38,203,907 parameters), we first pre-trained INTERACT by predicting DNAm levels of ∼25 million CpG sites of high coverage (> 50x) derived from a pseudo-bulk tissue comprised of all 4,137 nuclei from the same single-nucleus dataset. We then fine-tuned the pre-trained INTERACT model for each cell type by predicting the DNAm levels of 2.3 to 3.1 million CpG sites, chosen based on a coverage cutoff specific to each cell type (**Table S1**).

**Figure 1.**
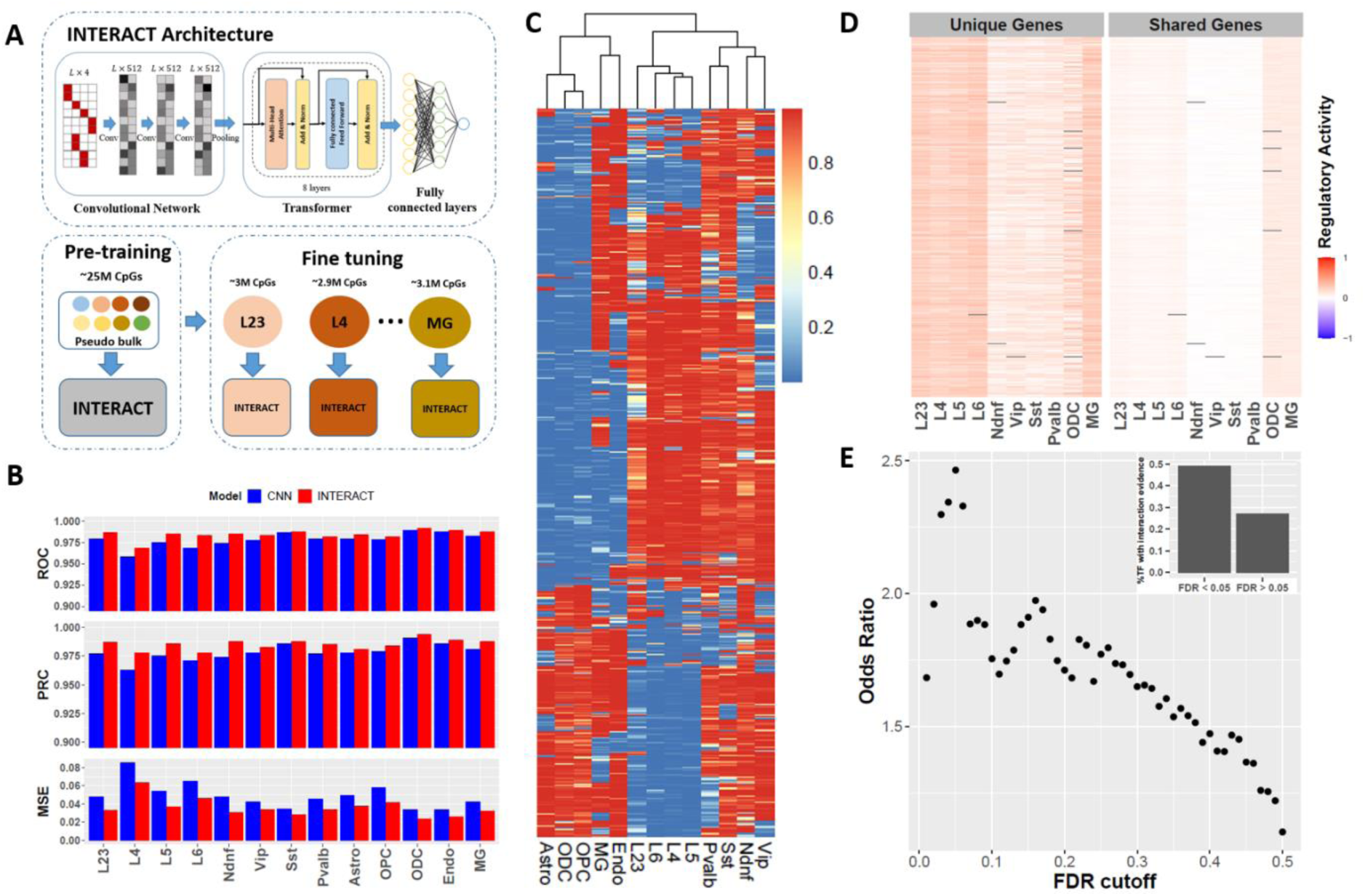
Prediction of cell type-specific DNAm levels from local DNA sequence. **A.** Illustration of INTERACT architecture and two-step training strategy (pre-training and fine- tuning). **B.** Comparison of model performance between INTERACT and CNN in predicting DNAm levels of independent CpG sites in each cell type. The performance was evaluated using three metrics: area under the Receiver Operating Characteristic curve (ROC), area under the Precision-Recall Curve (PRC), and mean squared error (MSE). ODC, oligodendrocyte. Astro, astrocyte. MG, microglia. OPC, oligodendrocyte progenitor cell. Endo, endothelial cell. L2/3, L4, L5 and L6 denote excitatory neuron subtypes in different cortical layers. Pvalb and Sst, medial ganglionic eminence-derived inhibitory subtypes. Ndnf and Vip, CGE-derived inhibitory subtypes. **C.** Hierarchical clustering of 13 brain cell types from predicted DNAm levels of independent CpG sites on chromosome 22 that were not used for model training and validation. **D.** Filters show higher regulatory activity for enhancers targeting cell type-specific genes compared to enhancers targeting shared genes. **E.** Enrichment of physical interaction evidence with DNA methylation or demethylation enzymes for TFs identified at varying FDR cutoffs. The upper corner highlights the proportions of TFs with physical interaction evidence in two groups, separated by an FDR cutoff of 0.05.

We evaluated the performance of each cell type-specific INTERACT model in predicting the DNAm levels of independent CpG sites not included in both pre-training and fine-tuning stages. We observed remarkable prediction performance across all cell type-specific models when measured by their abilities to distinguish methylation from unmethylation status of CpG sites, with an average area under the Receiver Operating Characteristic (ROC) curve of 0.984 and an average area under the Precision-Recall Curve (PRC) of 0.985. We compared the performance of INTERACT with a standard CNN model, and INTERACT consistently outperformed the CNN model (**Figure 1B**). The INTERACT models showed higher average ROC (0.984 for INTERACT vs. 0.978 for CNN), higher average PRC (0.985 for INTERACT vs. 0.978 for CNN), and lower average mean squared error (MSE) values (0.0358 for INTERACT vs. 0.0491 for CNN).

We further assessed whether cell type-specific INTERACT models capture cell type-specific information. To do this, we predicted DNAm levels of independent CpG sites on chromosome 22 for each cell type. We then selected the top 1000 variable CpG sites across 13 brain cell types and performed clustering analysis for the 13 brain cell types. We observed that predicted top variable CpG sites can clearly cluster cell types that align with their biological relationships (**Figure 1C**), suggesting that our cell type-specific models effectively learned cell type-specific information for predicting DNAm levels.

### Cell type-specific INTERACT models learn DNA motifs and transcription factors underlying cell type-specific DNAm profiles

To explore the mechanism underlying how trained cell type-specific INTERACT models learned cell type-specific information, we examined filters in the first convolutional layer of each model in their ability to activate the expression of cell type-specific genes. To achieve this goal, we defined a regulatory activity for each filter, quantifying the correlation between the activation strength of filters within enhancer regions and the expression levels of the genes targeted by these enhancers. A higher regulatory activity reflects a greater potential of the filter to activate genes targeted by enhancers. Our findings revealed that filters learned by each cell type-specific model tend to have a higher regulatory activity for enhancers targeting cell type-specific genes, compared to enhancers targeting shared genes between the corresponding cell type and at least one other cell type (**Figure 1D**). This analysis suggests that our cell type-specific models effectively learned DNA motifs that play crucial roles in cell type-specific gene regulatory circuits, thus enabling them to capture cell type-specific information.

To gain further biological insights from the trained cell type-specific INTERACT models, we also examined filters in the first convolutional layer to identify DNA motifs learned by these filters for each cell type. We then compared learned DNA motifs with known transcription factor (TF) binding motifs using the Tomtom motif comparison tool(*24*). In total, we identified 143 unique TFs (FDR < 0.05) whose DNA binding motifs matched the filter-learned DNA motifs and that were expressed in the corresponding cell type (**Data S1**). The number of detected TFs varied for each cell type-specific model, ranging from 45 for the excitatory neuron (L23) and microglia to 73 for the inhibitory neuron (Sst), with an average of 58 TFs for each cell type.

Among the 143 TFs, 48% (69) showed physical interaction evidence with enzymes involved in DNA methylation (DNMT1, DNMT3A, DNMT3B) or demethylation (TET1, TET2, TET3 and TDG). This represents a significant enrichment (OR = 2.5, p = 1.8 × 10^-6^) compared to a list of background TFs whose DNA binding motifs did not match by any filter-learned DNA motifs (835 TFs, FDR > 0.05), with only 24% of them showing such evidence. Furthermore, we explored the enrichment pattern of physical interaction evidence for TFs detected at varying FDR thresholds, and found an increasing trend of enrichment when FDR cutoffs ranged from 0.01 to 0.05, followed by a decrease in enrichment when FDR cutoffs went beyond 0.05 (**Figure 1E**). This enrichment pattern indicates an intriguing relationship between filter-indicated TFs and enzymes responsible for the biochemical processes of DNAm, suggesting that our cell type-specific models may learn TFs that regulate cell type-specific DNAm profiles.

### Predicted cell type-specific DNAm regulatory variants reflect cellular context

We performed *in silico* mutagenesis to predict variant effects on DNAm levels of CpGs in specific cell types based on the trained cell type-specific INTERACT models (**Figure 2A**). We then sought to evaluate whether predicted cell type-specific variant effects reflect the cellular context. Firstly, we hypothesized that if the predicted cell type-specific effects captured the cellular context, then variants would show similar effects within similar cell types. Indeed, we observed a stronger correlation pattern for the effects of variants predicted by models trained on similar cell types, and clustering of the pairwise correlation matrix clearly formed three distinct clusters: excitatory, inhibitory, and non-neuronal cell types (**Figure 2B**). Specifically, excitatory neuron subtypes (L2/3, L4, L5, and L6) were clustered together, while inhibitory neuron subtypes (Ndnf, Pvalb, Sst and Vip) formed a related but distinct cluster, and non-neuronal cell types were grouped together and separated from neuronal cell types, aligning with the biological relatedness of these cell types.

**Figure 2.**
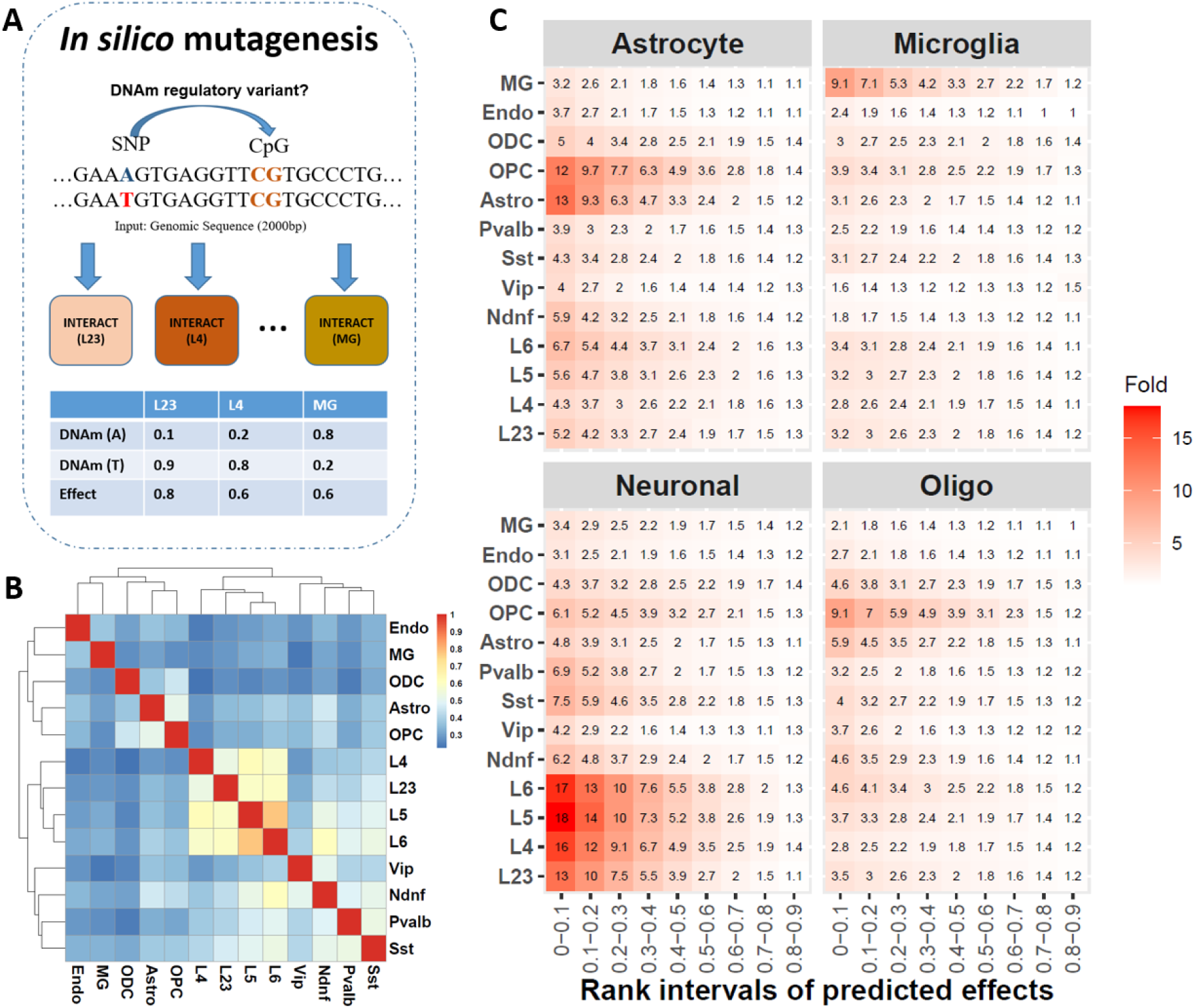
*In silico* discovery and characterization of cell type-specific DNAm regulatory variants. **A.** Schematic view of *in silico* discovery of cell type-specific DNAm regulatory variants. **B.** Clustering of 13 brain cell types by their predicted SNP effects on DNAm levels from each cell type-specific INTERACT model. **C.** Enrichment of active enhancers unique to each broad brain cell type (neuron, astrocyte, microglia, and oligodendrocyte (Oligo)) among variants ranked at different intervals by their predicted effects on DNAm levels in each cell type. Rank interval “0-0.1” represents variants of large effect and ranked in the top 10%. The color gradient represents enrichment fold of variants in each rank interval for their enrichment of enhancers compared to variants ranked in the bottom 10%.

Secondly, given the role of DNAm in gene regulation, we hypothesized that if the predicted variant effects on DNAm are cell type-specific, variants with higher effects would be enriched in active regulatory regions unique to the corresponding cell type. To test this, we overlapped variants with active enhancers (marked by histone modification H3K27ac outside of H3K4me3) and unique to each of the four broad cell types in the human brain (neuron, astrocyte, microglia, and ODC) determined based on a previous study(*25*) (**Figure 2C**). Our analysis confirmed that variants with higher effects in one cell type were more enriched in active enhancers in the corresponding or closely related cell types, as compared to variants with lower effects (ranked in the bottom 10%). These results provide further evidence that the predicted cell type-specific effects of variants capture the cellular context.

### Predicted cell type-specific DNAm regulatory variants were enriched for the heritability of brain-related traits in relevant cell types

We performed stratified LD-score regression (S-LDSC) to evaluate the contribution of predicted cell type-specific DNAm regulatory variants to the genetic components of 18 brain-related traits. We found that variants with large effect (ranked in the top 10%) in neuronal cell types were enriched for the heritability of a number of brain-related traits and disorders (**Figure 3**), aligning with the known neurobiology of these traits. For example, variants in both excitatory and inhibitory neurons showed strong enrichment for schizophrenia, bipolar disorder and depression. Among three neurological diseases we examined, we observed strong enrichment for epilepsy in both excitatory and inhibitory neurons, but not in non-neuronal cell types, consistent with the cell type-specificity analysis using gene expression signatures from a recent GWAS study(*26*). For Alzheimer’s disease (AD), we observed the strongest enrichment in microglia (fold = 5.2), but with only a trend for nominal significance (*p* = 0.071), probably due to the limited power of current AD GWAS. We also noted the enrichment of predicted DNAm regulatory variants in non-neuronal cell types for the heritability of multiple brain-related traits, though to a lesser extent compared to neuronal cell types. For example, variants in astrocytes were enriched for heritability of schizophrenia and bipolar disorder, while variants in astrocytes, microglia, ODC and endothelial cells showed enrichment for depression heritability, consistent with the emerging roles of non-neuronal cell types in psychiatric disorders(*27*).

**Figure 3.**
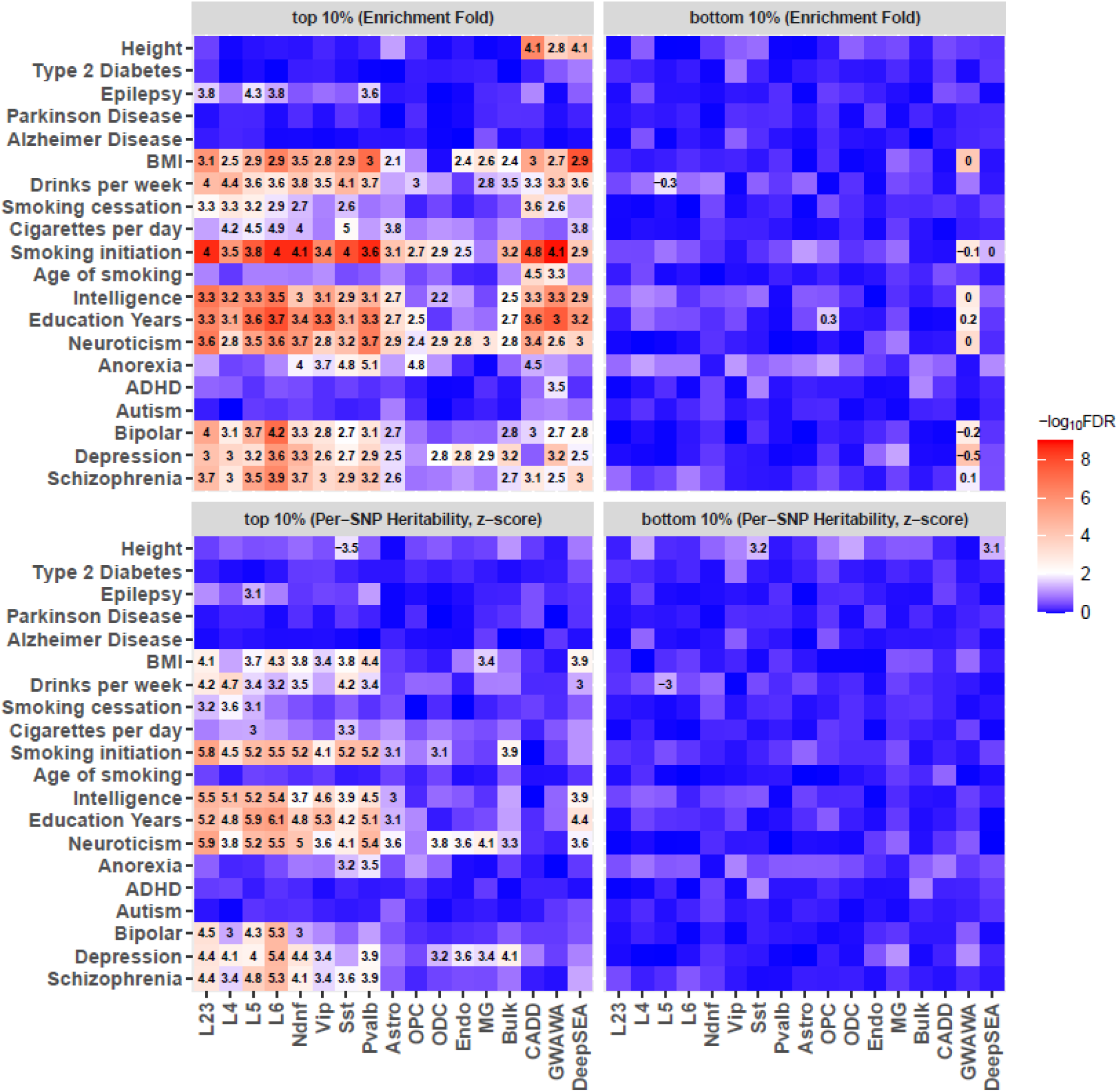
Comparison of variants ranked by different scoring systems for their contributions to heritability of brain-related traits. The left and right panel represents variants of high effect (in the top 10%) and low effect (in the bottom 10%), respectively, as predicted by the INTERACT model of each brain cell type or bulk brain sample (DLPFC) and three additional scoring systems (CADD, GWAWA and DeepSEA). The color gradient represents significance levels (FDR) for enriched heritability (upper panel) or z-score of per-SNP heritability (lower panel). The numbers within the squares of upper panel are enrichment fold that are significant after multiple testing correction (FDR < 0.05). The numbers within the squares of lower panel are z-score of per-SNP heritability that are significant after multiple testing correction (FDR < 0.05).

Additionally, we compared these findings with the top 10% ranked variants predicted by the INTERACT model trained on bulk brain sample in our previous study(*23*) (**Figure 3**). While we also observed a certain level of enrichment for brain-related traits from the bulk model, the strength of enrichment was generally lower compared to the INTERACT models trained on neuronal cell types. This comparison emphasizes that cellular context matters for predicting functional variants underlying brain-related traits. We also compared top-ranked variants from the cell type-specific INTERACT models with variants ranked by three other scoring systems, including CADD(*28*), GWAWA(*29*) and DeepSEA(*30*) (predicted effects on enhancer mark H3K27ac in frontal cortex) (**Figure 3**). While enrichment of heritability was also observed for brain-related traits in the variants ranked by these scores, their strength tended to be weaker compared to variants ranked by cell type-specific INTERACT models.

To further assess the impact of predicted DNAm regulatory variants on trait heritability, we employed another metric from S-LDSC: the z-score of per-SNP heritability. This metric allows us to discern the unique contributions of predicted variants to trait heritability while accounting for contributions from other functional annotations in the baseline model. We observed compelling statistical evidence (FDR < 0.05) supporting the involvement of top-ranked variants in neuronal cell types for 12 brain-related traits (**Figure 3**). However, when we examined the top-ranked variants in bulk brain samples, only three traits (depression, neuroticism and smoking initiation) showed significant z-score (FDR < 0.05), and no traits displayed significant z-score for variants ranked by CADD and GWAWA. While DeepSEA-ranked variants showed significant contributions to five brain-related traits (FDR < 0.05), their statistical evidence was weaker when compared to variants ranked by INTERACT models trained on neuronal cell types. This analysis indicates that the variants ranked by our cell type-specific models offer unique and significant contributions to the genetic components of brain-related traits.

To investigate the specificity of our findings in brain, we examined whether predicted cell type-specific DNAm regulatory variants showed enrichment for the heritability of human height and type 2 diabetes, serving as two negative controls. We did not observe significant enrichment for top-ranked variants in any cell types for these two traits, implying that the effects of these variants were relatively specific to brain-related traits. In contrast, top-ranked variants by the other three scoring systems (CADD, GWAWA and DeepSEA) showed strong enrichment for human height, suggesting their general roles in gene regulation rather than being specific to brain-related traits. Furthermore, we performed S-LDSC for variants with low effects (ranked in the bottom 10%) and found no evidence of heritability enrichment for brain-related traits in any cell types, suggesting that the observed enrichment for brain-related traits were specific to variants with large effects. Finally, we conducted S-LDSC for variants ranked at other different levels, and in general, we observed less evidence of their contributions to trait heritability for variants with weaker effects (**Data S2-3**).

### Cell type-specific INTERACT models enhance fine mapping for brain disorders

Given the roles of DNAm regulatory variants in brain-related traits, we reasoned that causal variants for brain disorders would have a stronger impact on DNAm levels in cell types relevant to the disorders. To test this, we conducted fine mapping for the GWAS risk loci of three major brain disorders (schizophrenia, depression and AD), and then compared variants with a higher chance of being causal (posterior inclusion probability (PIP) > 0.1 and p < 1 × 10^-6^) to control variants of less likely causal (PIP < 1 × 10^-4^ and p > 0.99), in terms of their predicted effects on DNAm levels and DNAm levels of their affected CpG sites in each brain cell type.

We observed that putative causal variants for schizophrenia tend to have a larger effect on DNAm levels than control variants in all neuronal cell types (except Sst) and astrocytes, with no significant difference in other glia cells (p > 0.05) (**Figure 4A**). Group difference was also noticed from INTERACT trained on bulk brain sample, though with weaker significance (p = 0.01) compared to INTERACT trained on neuronal cells, highlighting the importance of cellular context in understanding the functional impacts of risk variants. For depression, we observed group differences in all neuronal cells (except Vip) and three types of glia cells (astrocytes, OPC, and ODC). For AD, differences were observed in all excitatory neurons, one inhibitory neuron (Sst), and all glia cells (except ODC). Notably, difference in microglia was unique to AD among the three disorders, consistent with the distinct role of microglia in AD. Interestingly, the recent single-nucleus RNA-Seq study of AD detected the largest number of differentially expression genes in excitatory neurons and found depletion in the inhibitory neuron (Sst) in AD(*31*). This finding aligns with the neuronal cell types we observed, where AD putative causal variants show stronger effects. Additionally, we compared the DNAm levels of CpG sites impacted by the two groups of variants. We observed a trend of lower DNAm levels for putative causal variants across all three brain disorders (**Figure 4B**), suggesting that risk variants tend to impact genomic regions with a regulatory potential, as indicated by their low DNAm levels.

**Figure 4.**
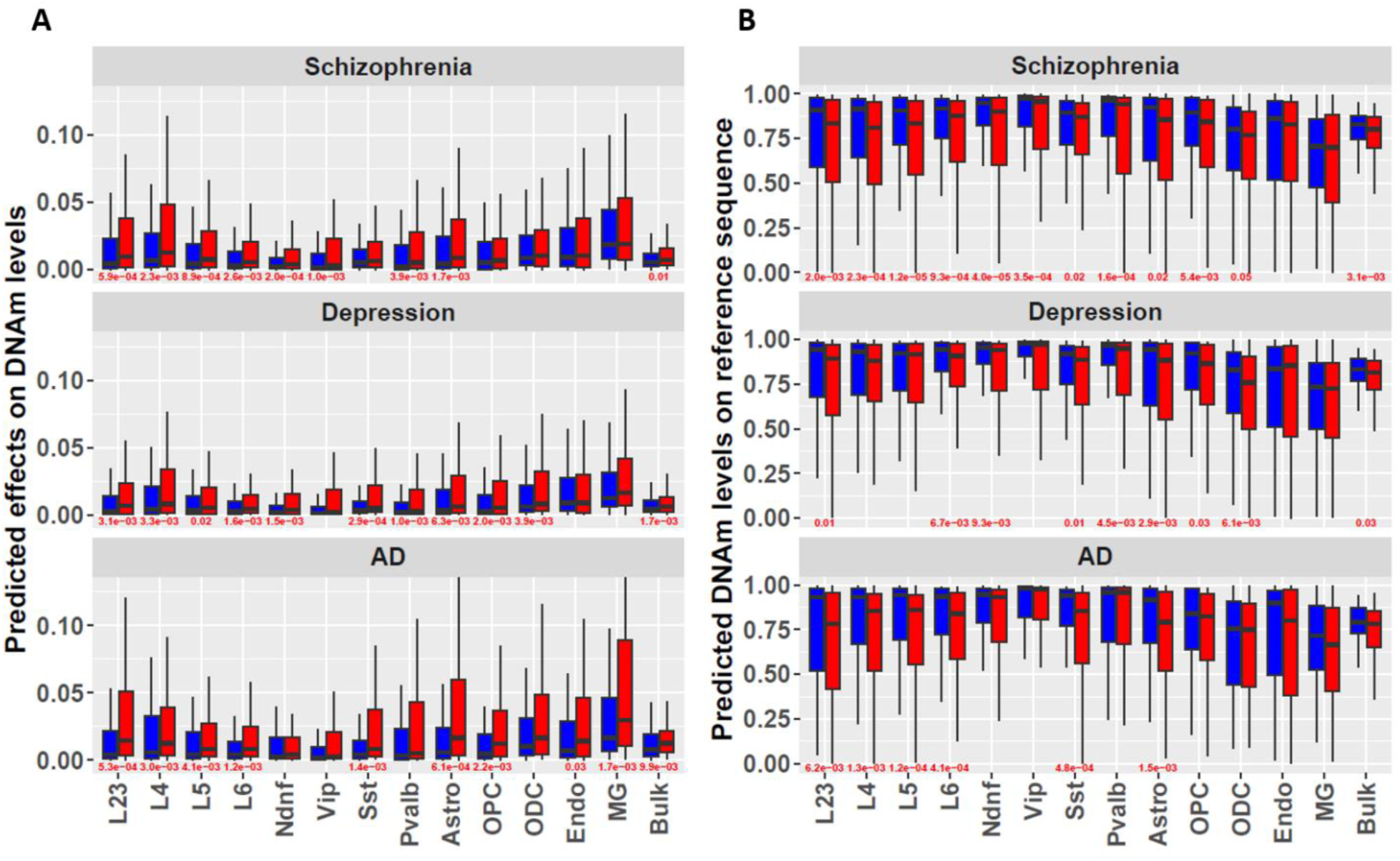
Predicted variant effects and DNAm levels of CpG site differentiate putative causal variants from control variants for three brain disorders. **A.** Fine-mapped putative causal variants show stronger impacts on DNAm levels in specific brain cell types than variants that are unlikely to be causal. **B.** Fine-mapped putative causal variants show lower DNAm levels for their impacted CpG sites in specific brain cell types than variants that are unlikely to be causal. Red numbers indicate significant group difference (p < 0.05, Wilcoxon rank-sum test).

Given our observation that putative causal variants have a stronger impact on DNAm levels and their affected CpG sites have lower DNA levels in specific cell types, we aimed to determine whether incorporating these predicted functional annotations could improve the fine mapping of risk loci underlying the three aforementioned brain disorders. To incorporate predicted functional annotations into fine mapping framework, we employed CARMA, a novel Bayesian approach that can jointly model GWAS summary statistics and functional annotations(*32*). We first performed fine mapping by including 26 functional annotations predicted from all of 13 cell type-specific models. Compared to fine mapping without annotations, we observed a much smaller number of SNPs included in the 99% credible sets from fine mapping with annotations (**Figure 5A**), and a larger number of risk loci counted by the maximum number of SNPs (up to 10) included in the 99% credible sets (**Figure 5B, 5C, 5D**). We provide details for the fine-mapped SNPs (PIP > 0.05 in 99% credible sets), their predicted functional annotations, and their assigned gene targets for each disorder in **Data S4-6**, offering potential candidate genes for further investigation.

**Figure 5.**
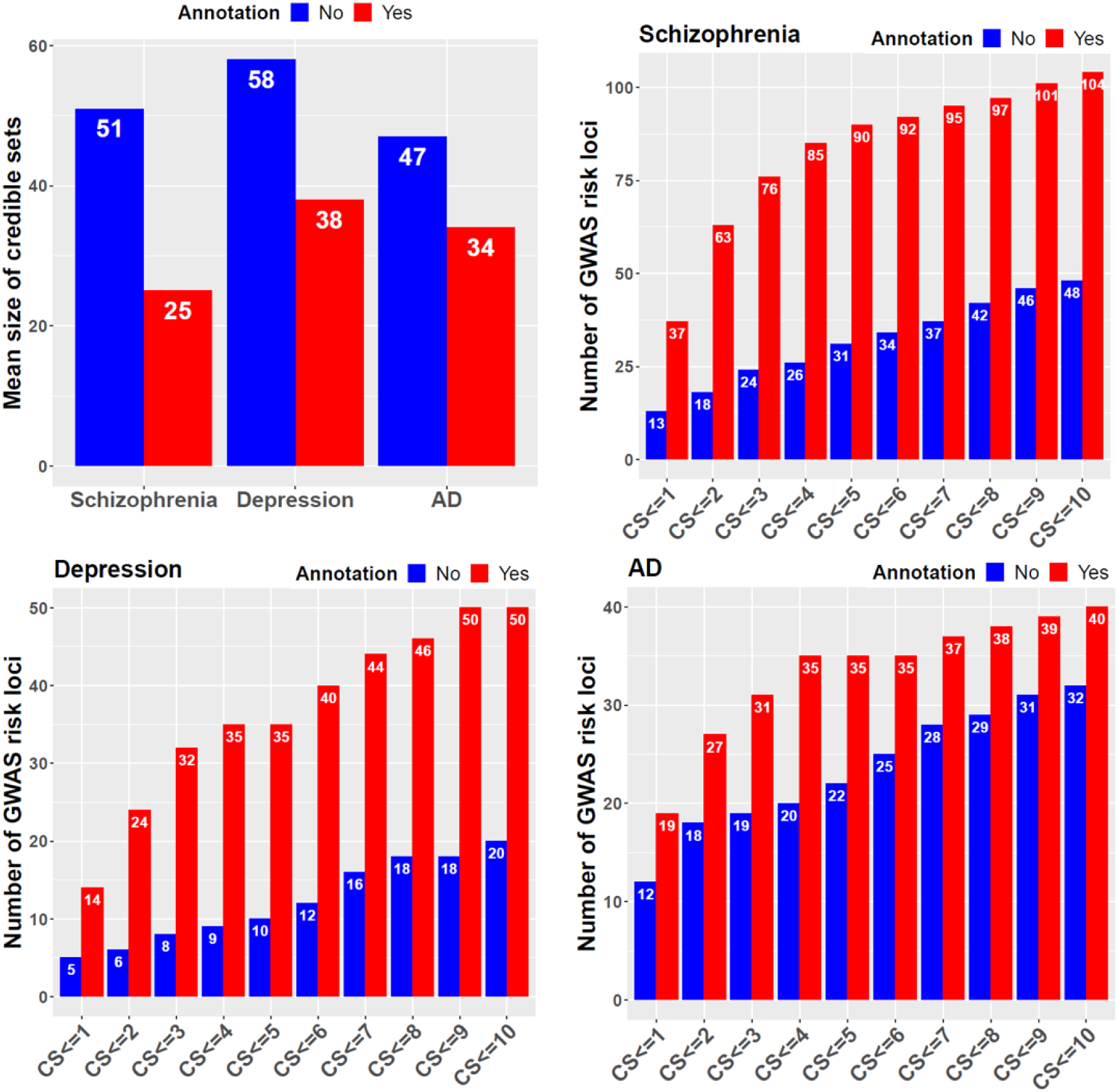
Functional annotations predicted by cell type-specific INTERACT models enhance fine mapping for three brain disorders. Comparison of the average size of credible sets between fine mapping with and without annotations (**top left**). Comparison of the number of risk loci counted by the maximum number of SNPs (up to 10) included in the 99% credible sets between fine mapping with and without annotations for schizophrenia (**top right**), depression (**bottom left**) and AD (**bottom right**)

Following the fine mapping strategy that incorporated annotations from all 13 brain cell types, we proceeded with additional fine mapping using annotations from refined cell types. These refined cell types encompassed four broad categories (excitatory neuron, inhibitory neuron, neuron, and glia), along with each individual cell type. Our objective was to pinpoint specific cell types that might yield a more robust fine mapping signal, offering insights into where the causal variants and their target genes are likely to exert their effects. Indeed, we observed many risk loci showing stronger PIP signals in certain cell types across the three disorders, whereas signals in other cell types were notably weaker or absent (**Data S7-9**). We illustrate one such risk loci for AD on chromosome 7, where fine mapping without annotations generated a credible set of six SNPs, with the highest PIP observed for rs74504435 (PIP = 0.37) (**Figure 6**). Notably, fine mapping with annotations from all cell types produced a credible set of only two SNPs, with the highest PIP=0.986 observed again for rs74504435. Interestingly, fine mapping using annotations from refined cell types clearly identified rs74504435 in glia (PIP = 0.981), specifically in astrocytes (PIP = 0.917). In contrast, among other refined cell types, the highest PIP observed for this SNP was only 0.429, found in one neuronal cell type (L6). Further investigation into the function of rs74504435 revealed that this SNP had the largest impact on a CpG site (chr7: 54949625), situated within an enhancer in astrocytes that exhibited Hi-C contact with the promoter of EGFR (epidermal growth factor receptor). Furthermore, the risk allele “A” of rs74504435 was predicted to decrease DNAm level of the CpG site (DNAm level = 0.19) compared to the reference allele “C” (DNAm level = 0.46). Given prior evidence that regulatory regions are associated with low DNAm levels, we hypothesize that the risk allele “A” could increase the enhancer activity and hence increase the expression level of *EGFR.* Intriguingly, previous studies have shown that over-expression of *EGFR* can lead to amyloid-β induced memory loss(*33*) and induce neuroinflammation and activate astrocytes(*34*). Additionally, inhibitors of EGFR have been explored as potential treatments for AD(*35*).

**Figure 6.**
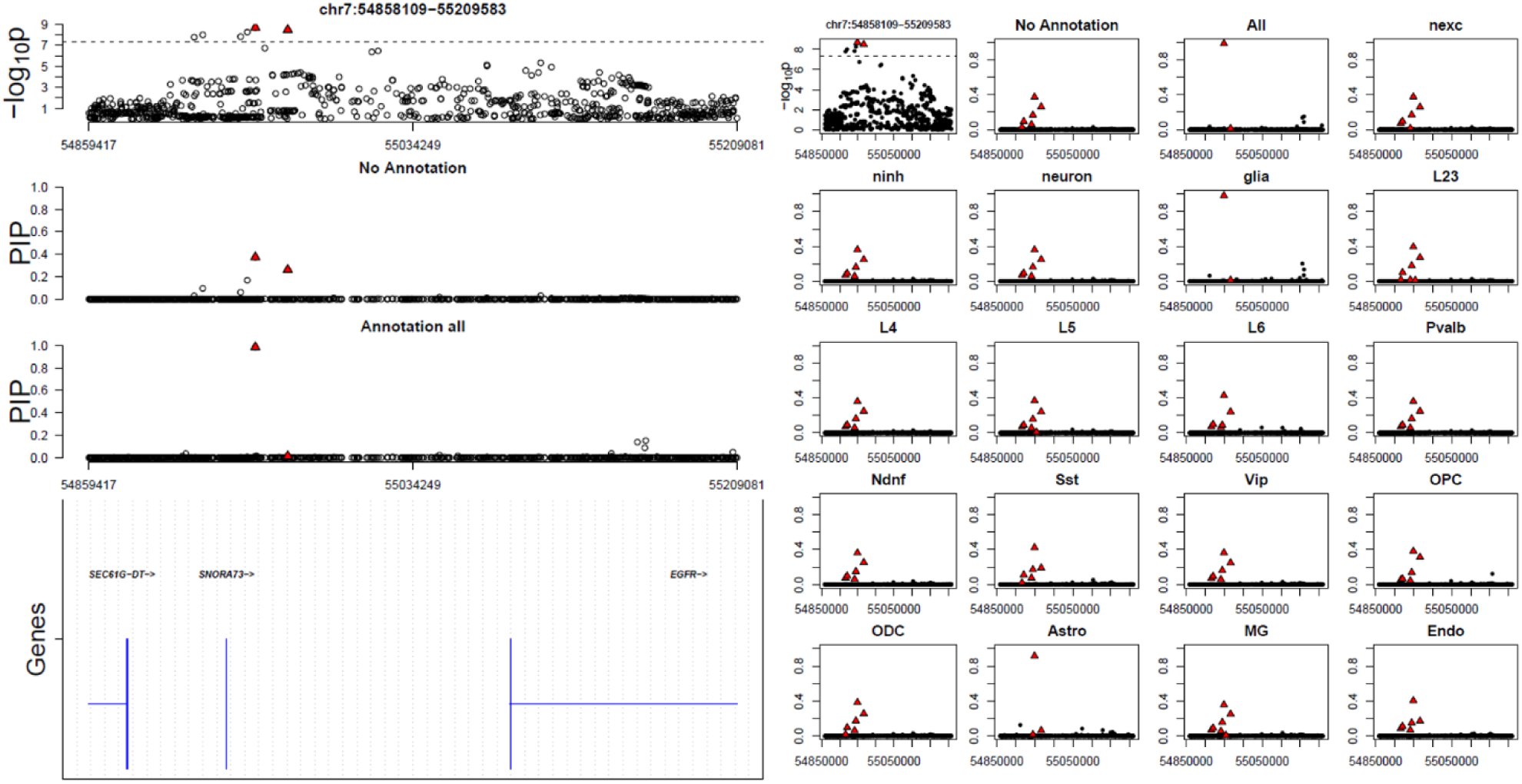
Fine-mapping of an AD GWAS risk locus identifies a causal gene in astrocyte. The left panel shows a regional plot for GWAS association signals, followed by a regional plot for posterior inclusion probability (PIP) computed by CARMA without functional annotations, a regional plot for PIP computed by CARMA with functional annotations from 13 brain cell types, and genes within the locus. Red triangles indicate fine-mapped SNPs within the 99% credible set obtained from fine mapping with all annotations. The right panel shows a regional plot of GWAS association signals for the same locus, followed by regional plots for PIP computed by CARMA without functional annotations, a series of regional plots for PIP computed by CARMA with functional annotations from all cell types, four broad cell types (nexc, ninh, neuron, glia), and 13 individual cell types. nexc: excitatory neuron; ninh: inhibitory neuron; Red triangles indicate fine-mapped SNPs within the 99% credible set from fine mapping with each type of annotation; Red triangles in GWAS regional association plot indicate fine-mapped SNPs within the 99% credible set obtained from fine mapping with all annotations.

## Discussion

In our previous study, we introduced the INTERACT model, designed to predict DNAm regulatory variants in bulk brain samples(*23*). This current study extends the INTERACT model to identify DNAm regulatory variants in specific brain cell types utilizing existing single-nucleus DNAm data from the human brain. Our study reveals the inherent power of DNA sequences to encode cell type-specific DNAm patterns and demonstrate the ability of our cell type-specific INTERACT models to reveal DNA motifs and TFs underlying cell type-specific DNAm profiles. We show that our cell type-specific INTERACT models predict DNAm regulatory variants that capture cellular context and are enriched for the heritability of many brain-related traits in relevant cell types. Importantly, we demonstrate that the inclusion of predicted SNP effects and DNAm levels enhances the fine mapping of risk loci for three major brain disorders and pinpoint particular cellular types where the fine-mapped risk variants may exert their effects. Our findings highlight the power of deep learning in identifying functional regulatory variants in specific cell types, which will further our understanding of the genetic underpinnings of complex traits.

This work should be viewed in light of several limitations. Firstly, our model relies on input DNA sequences from the reference genome, which may not precisely align with the DNA sequences in the samples we used for training the model. The performance of the model could be further improved in future research by utilizing DNA sequences and DNAm data from the same individuals. Secondly, our study was limited by the number of nuclei available in the single- nucleus DNAm data we utilized, allowing us to build cell type-specific models for only 13 brain cell types. This hinders our ability to identify DNAm regulatory variants in more refined cell types. This limitation could be addressed by incorporating larger-scale single-nucleus DNAm datasets that cover a wider range of cell types. Thirdly, given the high sparsity of single-nucleus DNAm profile and the need for large training samples, we assumed that the DNAm status of a CpG site is consistent across nuclei of the same type, which allowed us to aggregate training samples of CpG sites that are fully methylated or unmethylated across nuclei of the same type. However, this assumption may not always hold true, particularly for nuclei within a broad cell type. This limitation could be addressed by increasing sequencing coverage in individual nucleus or collecting a larger number of nuclei for sequencing, which may lead to a more accurate measurement of DNAm levels of CpG sites for each cell type, including those of intermediate DNAm levels. Lastly, variants derived from our model can only assist in uncovering risk variants that act through regulation of DNAm, but it is unable to reveal risk variants that operate through mechanisms independent of DNAm regulation. Our model has potential to be extended to identify regulatory variants for other types of molecular traits, such as gene expression and histone modification in specific cell types.

## Methods

### Training datasets

We trained the cell type-specific INTERACT models using an existing single-nucleus dataset generated by the single-nucleus methyl-3C sequencing technique (sn-m3C-seq)(*16*), which allows simultaneous capture of DNAm and chromatin contact information from individual nuclei. This dataset includes 4,137 nuclei derived from the human prefrontal cortex, which were clustered into 13 cell types. These cell types include four excitatory neuron subtypes characteristic of different cortical lamina (L2/3, L4, L5, and L6), four inhibitory neuron subtypes (Ndnf, Vip, Pvalb, and Sst), and five non-neuronal subtypes (astrocyte, oligodendrocyte (ODC), oligodendrocyte progenitor cell (OPC), microglia, and endothelial cell). To build the training dataset for each cell type, we utilized pseudo-bulk tissue derived from the nucleus of the corresponding cell type. We carefully selected CpG sites that were either fully methylated or fully unmethylated as the training samples for each cell type. Our rationale for this approach was two-fold: First, we assumed that the DNAm status at a specific CpG site tends to be consistent across nuclei of the same cell type. Second, due to the high sparsity of single-nucleus DNAm profiles, it was challenging to collect a large number of CpG sites with sufficient read coverage (> 50) for reliable estimation of intermediate DNAm levels, especially for neuronal cell types characterized by very limited number of nuclei in this dataset (**Table S2**). Consequently, our training dataset for each cell type included only CpG sites that were either fully methylated or fully unmethylated. To maintain data quality while ensuring an adequate training sample size, we implemented a manually adjusted coverage cutoff for each cell type. As a result, the cell type-specific training datasets contained between 2.3 and 3.1 million CpG sites. **Table S1** provides details on the coverage threshold and training sample size for each cell type. The pre-training dataset for INTERACT comprises approximately 25 million CpG sites obtained from a pseudo-bulk tissue derived from all 4,137 nuclei mentioned earlier. DNAm levels were determined by calculating the ratio of methylated reads to the total number of reads across all nuclei. To ensure high data quality for our pre-training process, we only included CpG sites covered by more than 50 reads across all nuclei.

For both the pre-training and cell type-specific training datasets, we divided CpG sites into three subsets by chromosomes for model training, validation, and evaluation. The training set consisted of CpG sites on chromosomes 1 to 20, while CpG sites on chromosome 21 were used as the validation set for model tuning, and CpG sites on chromosome 22 were used as the independent testing set to evaluate the model prediction performance.

### INTERACT architecture

INTERACT contains three main modules: CNN, the encoder module of transformer, and the fully connected network. The input to the INTERACT model is a one-hot encoded DNA sequence of 2 kb, and the DNAm level of the CpG site centered in the DNA sequence as output. Our choice of 2 kb input DNA sequences was supported by our previous study that investigated the performance of INTERACT for input DNA sequences of different lengths (1 kb, 2 kb, 3 kb, and 4 kb). The architecture of our model is detailed in **Table S3**, and the code is publicly available on our GitHub repository. Below are descriptions of each module.

The CNN module includes three convolution layers, each containing 512 kernels of length 10. Each convolutional layer is activated by a rectified linear unit (ReLU) function and is subsequently followed by a normalization layer. After the three convolution layers, a max-pooling layer is employed to capture the most significant features from the output of convolutional layers. To enhance the speed and stability of training, a batch normalization layer is utilized after the max-pooling layer. Additionally, a dropout layer with a rate of 0.5 is used following the batch normalization layer to prevent overfitting.

The transformer module receives the features learned from the CNN module. This module contains a stack of eight identical layers, each of which includes two sublayers. The first sublayer is a multi-head self-attention layer, and the second sublayer is a simple, position-wise fully connected feed-forward network. Each sublayer is followed by a normalization layer to improve the speed and stability of training, and a dropout layer with a rate of 0.1 to prevent overfitting. In addition, there is a residual connection around each sublayer.

The fully connected network includes a single hidden layer with 512 units, an output layer, and a dropout layer with a drop rate of 0.1 to prevent overfitting. The sigmoid function is used after the output layer to scale the predicted values into a range between 0 and 1. The output layer includes one unit that corresponds to the DNAm levels.

### Pre-training and fine-tuning

The INTERACT model was designed with approximately 38 million parameters. It is challenging to train a robust cell type-specific INTERACT model due to the limited number of training samples collected for each cell type-specific model. To overcome this challenge, we employed a two-step training strategy (**Figure 1A**). First, we pre-trained the INTERACT model by predicting DNAm levels for approximately 25 million CpGs, through aggregating DNAm data across 4,137 nuclei into a single pseudo-bulk DNAm dataset. DNAm levels of a specific CpG site were calculated as the ratio of methylated reads to the total number of reads across all nuclei. To ensure high-quality of the pre-training samples, we only used CpG sites covered by more than 50 reads. This pre-training phase allowed the model to learn informative features associated with DNAm levels that are shared across cell types. In the second step, we fine-tuned the pre-trained model by predicting the DNAm levels of the training samples collected for each cell type, allowing the model to learn cell type-specific features underlying DNAm levels.

### Comparison with CNN model

We compared our fine-tuned cell type-specific INTERACT model with a standard CNN model for their performance in predicting DNAm levels in each cell type. The CNN model shared the same structure as the CNN module within the INTERACT architecture, followed by the same fully connected network module as described above.

### DNA motif analysis

We defined a score of regulatory activity for each filter, which quantifies the correlation between the activation strength of filters within enhancers and the expression levels of the genes targeted by these enhancers. The activation strength of filters is a measure of the sequence similarity between the filter and the enhancer, with a higher activation value indicating a greater similarity of DNA sequences. A higher regulatory activity indicates a greater potential of the filter to activate genes targeted by enhancers. We calculated and compared two regulatory activity scores for each filter: one for enhancers targeting cell type-specific genes and the other for enhancers targeting shared genes between the corresponding cell type and at least one other cell type. To define enhancers and cell type-specific genes or shared genes for our study, we relied on enhancers and active promoters defined in a previous study using two histone marks (H3K4me3 and H3K27ac) for four broad cell types (neuron, microglia, ODC, and astrocyte)(*25*).

Specifically, cell type-specific genes were defined as those with an active promoter (defined by both H3K4me3 and H3K27ac) in the corresponding cell type but not in other broad cell types. Shared genes were defined as those with an active promoter in the corresponding cell type and at least one other broad cell type. We linked enhancers to their target genes based on Hi-C contact data collected for three broad cell types (neuron, ODC and astrocytes) from the same prior study(*25*), with additional supplement for the three broad cell types using cell-type specific chromatin loops called from the same nucleus in our training data. To obtain gene expression levels in specific brain cell types, we generated pseudo-bulk data for each cell type using single-nucleus transcriptomes dataset across six human cortical areas and associated cell type annotations downloaded from the ALLEN BRAIN data portal. Gene expression levels for each gene in each cell type were determined by aggregating the total number of reads assigned to the gene across all cells in the pseudo-bulk sample, which was then normalized to one million reads followed by log2 transformation. We calculated regulatory activity for filters from models trained by individual neuronal cell types (L23, L4, L5, L6, Ndnf, Vip, Sst, Pvalb) based on Hi-C contact in the broad neuron cell type. Similarly, we calculated regulatory activity for filters from models trained by ODC and microglia based on Hi-C contact in the broad ODC and microglia, respectively. We did not calculate regulatory activity for filters from models trained by astrocytes due to the limited number of of Hi-C contacts for this cell type.

We further examined filters in the first convolutional layer of each cell type-specific INTERACT model to identify DNA motifs associated with DNAm levels in each cell type. Following the method described previously(*36*), DNA motifs were discovered for each filter by a subset of sequences where the filter produces an activation value higher than half of the maximum activation value across all subsets of sequences the filer has scanned. Selected subsets of sequences were then aligned to generate a position weight matrix, which was further matched to annotated TF binding DNA motifs in the Homo sapiens CIS-BP database using the Tomtom v4.10.1 motif comparison tool(*24*). We considered matches at FDR < 0.05 significant for each filter. To assess the potential involvement of identified TFs in relation to DNA methylation levels, we investigated their physical interactions with enzymes involved in DNA methylation (DNMT1, DNMT3A, DNMT3B) or demethylation (TET1, TET2, TET3, and TDG)(*37*). We collected physical interaction evidence with DNMT3A and DNMT3B from a previous study(*38*) and from STRING database(*39*).

### *In silico* mutagenesis

We performed *in silico* mutagenesis to identify functional genetic variations impacting DNAm levels in each cell type (**Figure 2B**). Briefly, we first introduced a variant allele in a given DNA sequence of 1 kb flanking a CpG site. Each sequence (with or without the introduced variant allele) was then passed to each trained cell type-specific INTERACT model to get the predicted DNAm level of the CpG site centered within each sequence. The impact of the introduced variant on DNAm level of the CpG site was estimated by the difference of the predicted DNAm levels of the CpG site within the two sequences. We did this in silico mutagenesis for 9,042,066 SNPs (minor allele frequency > 0.005) observed in the European ancestry samples of 1000 Genomes and located within a 1 kb window of 25,564,506 autosomal CpG sites. This resulted in estimated effects on DNAm levels for 182,211,784 SNP–CpG pairs from each cell type-specific INTERACT model. SNPs were ranked in descending order by their maximum absolute values of predicted effects for all of their paired CpG sites.

To evaluate the contextual relevance of predicted brain cell type-specific regulatory variants, we investigated their enrichment for active regulatory regions unique to each major brain cell type, including neurons, microglia, astrocytes, and ODC. Active regulatory regions unique to each cell type were determined based on a prior study(*25*) and defined by the presence of active enhancers (H3K27ac peaks that were outside of H3K4me3 peaks) in each respective cell type but absent in the other three cell types.

### Stratified LD score regression

We performed stratified LD score regression (S-LDSC)(*40*) to evaluate the enrichment of heritability of brain-related traits for variants ranked at different intervals by their impacts on DNAm levels predicted by each cell type-specific INTERACT model. We also included two non-brain traits, human height and type 2 diabetes, as two negative controls to examine whether our findings are specific to brain-related traits. We downloaded GWAS summary statistics of each trait from the sources listed in **Data S10**. Following recommendations from the LDSC resource website (https://alkesgroup.broadinstitute.org/LDSCORE), S-LDSC was run for each list of variants with the baseline LD model v2.2 that included 97 annotations to control for the LD between variants with other functional annotations in the genome. We used HapMap Project Phase 3 SNPs as regression SNPs, and 1000 Genomes SNPs of European ancestry samples as reference SNPs, which were all downloaded from the LDSC resource website. To evaluate the unique contribution of predicted regulatory variants to trait heritability, we also utilized another metric from S-LDSC: the z-score of per-SNP heritability. This metric allows us to discern the unique contributions of candidate annotations while accounting for contributions from other functional annotations in the baseline model. The p-values are derived from the z-score assuming a normal distribution and FDR was computed from the p-values based on Benjamini & Hochberg procedure using R function.

To compare the performance of our cell type-specific models with other scoring systems in their abilities to predict functional variants enriched for trait heritability, we also performed S-LDSC for variants ranked by three other scoring systems: CADD(*28*), GWAWA(*29*) and DeepSEA(*30*). The CADD score measures the deleteriousness of variants and is derived from a support vector machine trained to distinguish variants that have survived natural selection from simulated mutations that are enriched for deleterious variants, utilizing features from diverse genomic annotations. GWAWA scores are derived from a random forest model trained to discriminate curated pathogenic variants from benign variants by integrating various genomic annotations. DeepSEA is another deep learning-based approach that employs a CNN model to estimate noncoding variant effects on chromatin. We considered only DeepSEA-predicted effects on the enhancer mark H3K27ac in the frontal cortex, as it represents the most relevant genomic element and tissue context for brain-related traits.

### Fine-mapping GWAS risk loci

We used PLINK to clump significant SNPs from GWAS summary statistics for schizophrenia(*41*) depression(*42*) and AD(*43*) of European ancestry samples into independent risk loci on autosomes. To achieve this, we first obtained index SNPs (*p* < 5 × 10^−8^) that were LD-independent and had r^2^ < 0.1 within a 3-Mb window. Next, we defined the risk loci for each index SNP as 50-kb upstream of the leftmost and 50-kb downstream of the rightmost SNPs that were within a 3-Mb window and had an r^2^ > 0.2 with the index SNP. We then merged risk loci that were within 50-kb, resulting in 177, 100 and 54 loci for schizophrenia, depression and AD, respectively.

We fine-mapped GWAS risk loci for each disorder using CARMA(*32*), a novel Bayesian fine mapping approach that jointly model GWAS summary statistics and functional annotations while accounting for discrepancies between summary statistics and LD from reference panels. We excluded risk loci within the major histocompatibility complex region (chr 6: 25–36 Mb) from this analysis due to extensive LD structure of this region. We also excluded risk loci on chromosome 19 (44519920:46477842) for AD from fine mapping due to long-range LD in this region and extreme values of the summary statistics as did in the previous study. LD was estimated from 50,000 randomly selected samples of European ancestry from the UKBB GWAS dataset. We employed two fine-mapping strategies: one without functional annotations and the other incorporating functional annotations from all cell types or various subsets of cell types. These functional annotations are variant effects on DNAm levels of CpG sites and the DNAm levels of the CpG sites (from reference DNA sequence) predicted by each cell type-specific INTERACT model. When a SNP was predicted for its effects on multiple CpG sites, we chose the CpG site for which the SNP has the largest predicted effect.

### Assign fine-mapped SNPs to target genes

We first assigned SNPs within 99% credible sets to putative causal genes if the genes had transcripts whose promoters overlapped with fine-mapped risk variants. Promoters were defined by 1-kb upstream and 1-kb downstream of the transcript start site, based on the GENCODE version 42 basic gene annotations. Putative causal genes were also assigned if promoters of their transcripts are distally contacted by fine-mapped SNPs in specific cell type, utilizing cell-type specific chromatin loops called from the same set of nuclei used for model training. Specifically, the cell-type specific chromatin loops were called at a 10-kb resolution by the SnapHiC method(*44*) from the Hi-C component of the same set of nuclei described in the training datasets section. Details for the SnapHiC method and identification of chromatin loops in these nuclei were described in the prior report(*44*). To link candidate SNPs to distal promoters, we extended each SNP by 1-kb in both directions to define a risk variant-region. If one of the chromatin loop bins intersected the risk variant-region and another bin overlapped a promoter of a transcript, the promoter was then distally linked with the SNP. For fine-mapped SNPs that did not overlap or distally contact any promoters, we assigned them to the nearest genes.

## Supporting information

Supplemental_DataS10

Supplemental_DataS1

Supplemental_DataS2

Supplemental_DataS3

Supplemental_DataS4

Supplemental_DataS5

Supplemental_DataS6

## Acknowledgments

We acknowledge the Psychiatric Genomics Consortium, UK BioBank, GSCAN (the GWAS & Sequencing Consortium of Alcohol and Nicotine), the International Parkinson’s Disease Genomics Consortium, and the CTGlab (the Complex Trait Genetics lab at the VU University Amsterdam the Amsterdam University Medical Centre) for making their GWAS results publicly available. We would like to thank the research participants and employees of 23andMe, inc. for making this work possible.

## Author contributions

Conceptualization: JZ, DRW, SH

Methodology: JZ, SH

Investigation: JZ, SH

Visualization: JZ, SH Supervision: DRW, SH

Writing—original draft: JZ, DRW, SH Writing—review & editing: JZ, DRW, SH

## Data and materials availability

Data S7-9 for fine mapping GWAS risk loci for schizophrenia, depression and AD

https://doi.org/10.6084/m9.figshare.24319177.v1

