## Supplementary_materials for "Deep learning predicts DNA methylation regulatory variants in specific brain cell types and enhances fine mapping for brain disorders"

Jiyun Zhou *et al.*

**This PDF file includes:**

Tables S1 to S3

**Other Supplementary Materials for this manuscript include the following:**

Data S1 to S10

| **Cell type** | **# of nuclei** | **Coverage cutoff (methylated)** | **Coverage cutoff (unmethylated)** | **# of CpGs (methylated)** | **# of CpGs (unmethylated)** | **Total # of CpGs** |
| --- | --- | --- | --- | --- | --- | --- |
| L23 | 551 | 30 | 30 | 1879748 | 1134274 | 3,014,022 |
| L4 | 131 | 10 | 8 | 1596875 | 1301346 | 2,898,221 |
| L5 | 180 | 12 | 10 | 1839440 | 1296925 | 3,136,365 |
| L6 | 86 | 7 | 5 | 1659919 | 1302156 | 2,962,075 |
| Ndnf | 144 | 11 | 8 | 1435657 | 938355 | 2,374,012 |
| Pvalb | 134 | 10 | 8 | 1829959 | 1158937 | 2,988,896 |
| Sst | 217 | 16 | 13 | 1493491 | 1223687 | 2,717,178 |
| Vip | 171 | 13 | 11 | 1640193 | 1022991 | 2,663,184 |
| Astro | 449 | 30 | 30 | 1639246 | 1143454 | 2,782,700 |
| MG | 422 | 25 | 28 | 1981637 | 1106193 | 3,087,830 |
| ODC | 1244 | 60 | 60 | 1637972 | 1495899 | 3,133,871 |
| OPC | 203 | 15 | 12 | 1610724 | 1343233 | 2,953,957 |
| Endo | 205 | 14 | 13 | 1587350 | 1135149 | 2,722,499 |

**Table S1. Training samples for each cell type**. Coverage refers to the number of reads supporting either fully methylation or fully unmethylation across all nuclei of the same cell type

| **Cell type** | **# of nuclei** | **# of CpG sites (>30 reads)** | **# of CpG sites (>50 reads)** |
| --- | --- | --- | --- |
| L23 | 551 | 20322957 | 9323696 |
| L4 | 131 | 60199 | 21342 |
| L5 | 180 | 273751 | 31984 |
| L6 | 86 | 24812 | 12099 |
| Ndnf | 144 | 86476 | 23823 |
| Pvalb | 134 | 65577 | 21774 |
| Sst | 217 | 2096153 | 53075 |
| Vip | 171 | 407726 | 34907 |
| Astro | 449 | 19986213 | 8766345 |
| MG | 422 | 19861138 | 7908566 |
| ODC | 1244 | 25632294 | 24914907 |
| OPC | 203 | 2068845 | 53075 |
| Endo | 205 | 1786383 | 50399 |

**Table S2. Count of CpG sites by two cutoffs of coverage in pseudo-bulk of each cell type**

| **Parameters** | **Pretraining** | **Fine-tuning** |
| --- | --- | --- |
| Number of layers in CNN module | 3 | 3 |
| Number of kernels in each CNN layer | 512 | 512 |
| Kernel size for each CNN layer | 10 | 10 |
| Kernel size for max-pooling layer | 20 | 20 |
| Step size for max-pooling layer | 20 | 20 |
| Dropout rate for dropout layer | 0.2 | 0.2 |
| Number of attention layers in Transformer module | 8 | 8 |
| Number of attention headers | 8 | 8 |
| Input size | 512 | 512 |
| Hidden size | 512 | 512 |
| Intermediate size | 2048 | 2048 |
| Output size | 512 | 512 |
| Dropout rate | 0.1 | 0.1 |
| Number hidden layer in fully connected network | 1 | 1 |
| Input size for fully connected network | 512 | 512 |
| Hidden size for fully connected network | 512 | 512 |
| Number of tasks (output size for fully connected network) | 1 | 1 |
| Initial learning rate | 0.000176 | 0.000176 |
| Number of learning rate warmup steps | 10000 | 10000 |

**Table S3. Hyperparameters for INTERACT models**

**Data S1. (separate file)**

Details of Tomtom motif comparison results for DNA motifs activated by filters in each cell type-specific model

Data S2. (separate file)

**Comparison of variants ranked by different scoring systems for their contribution to heritability of brain-related traits.** The left and right panel represents variants of higher effect and lower effect, respectively, as predicted by the INTERACT model of each brain cell type or bulk brain sample (DLPFC) and three additional scoring systems (CADD, GWAWA and DeepSEA). The color gradient represents significance levels (FDR) for enriched heritability (upper panel) or z-score of per-SNP heritability (lower panel). The numbers within the squares of upper panel represent enrichment or depletion fold that are significant after multiple testing correction (FDR < 0.05). The numbers within the squares of lower panel are z-score of per-SNP heritability that are significant after multiple testing correction (FDR < 0.05).

**Data S3. (separate file)**

Details of S-LDSC results for variants ranked by different systems

**Data S4. (separate file)**

Fine-mapped SNPs, predicted functional annotations and their targets for schizophrenia risk loci

**Data S5. (separate file)**

Fine-mapped SNPs, predicted functional annotations and their targets for depression risk loci

**Data S6. (separate file)**

Fine-mapped SNPs, predicted functional annotations and their targets for Alzheimer’s disease risk loci

**Data S10. (separate file)**

GWAS summary statistics for heritability enrichment analysis
