## Supplementary figures and images for "Deep learning predicts DNA methylation regulatory variants in specific brain cell types and enhances fine mapping for brain disorders"

### Supplemental_DataS2

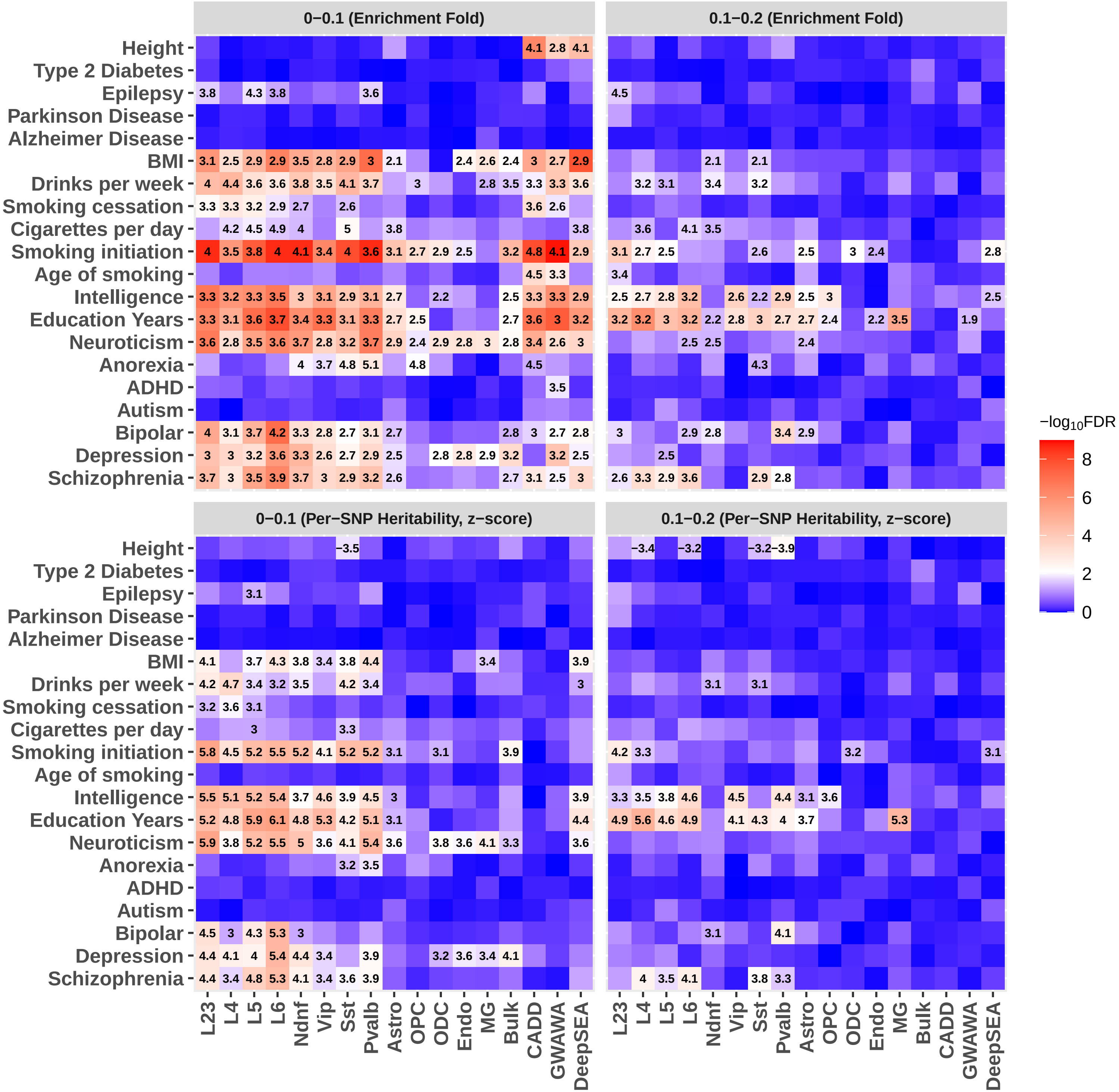

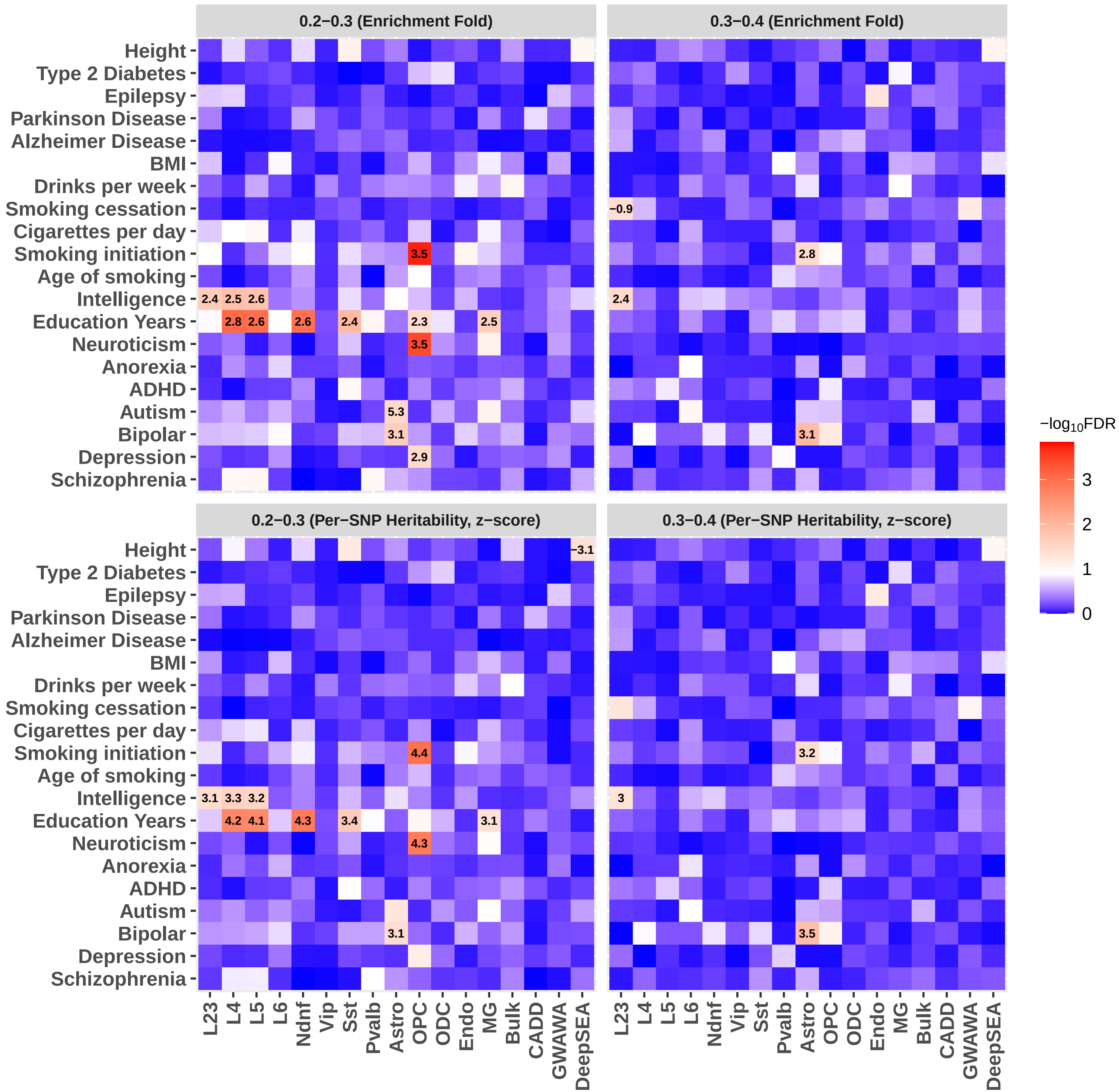

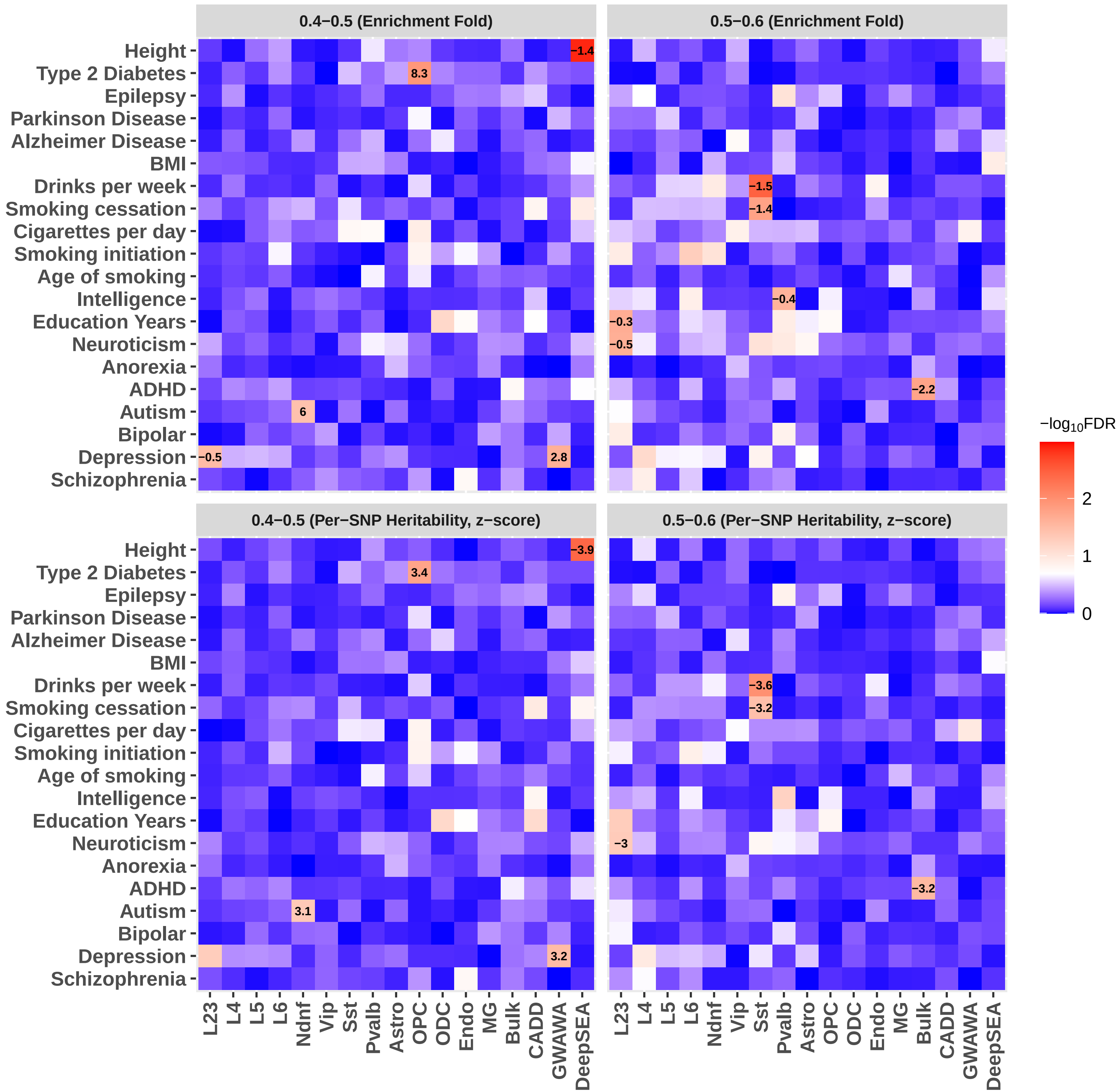

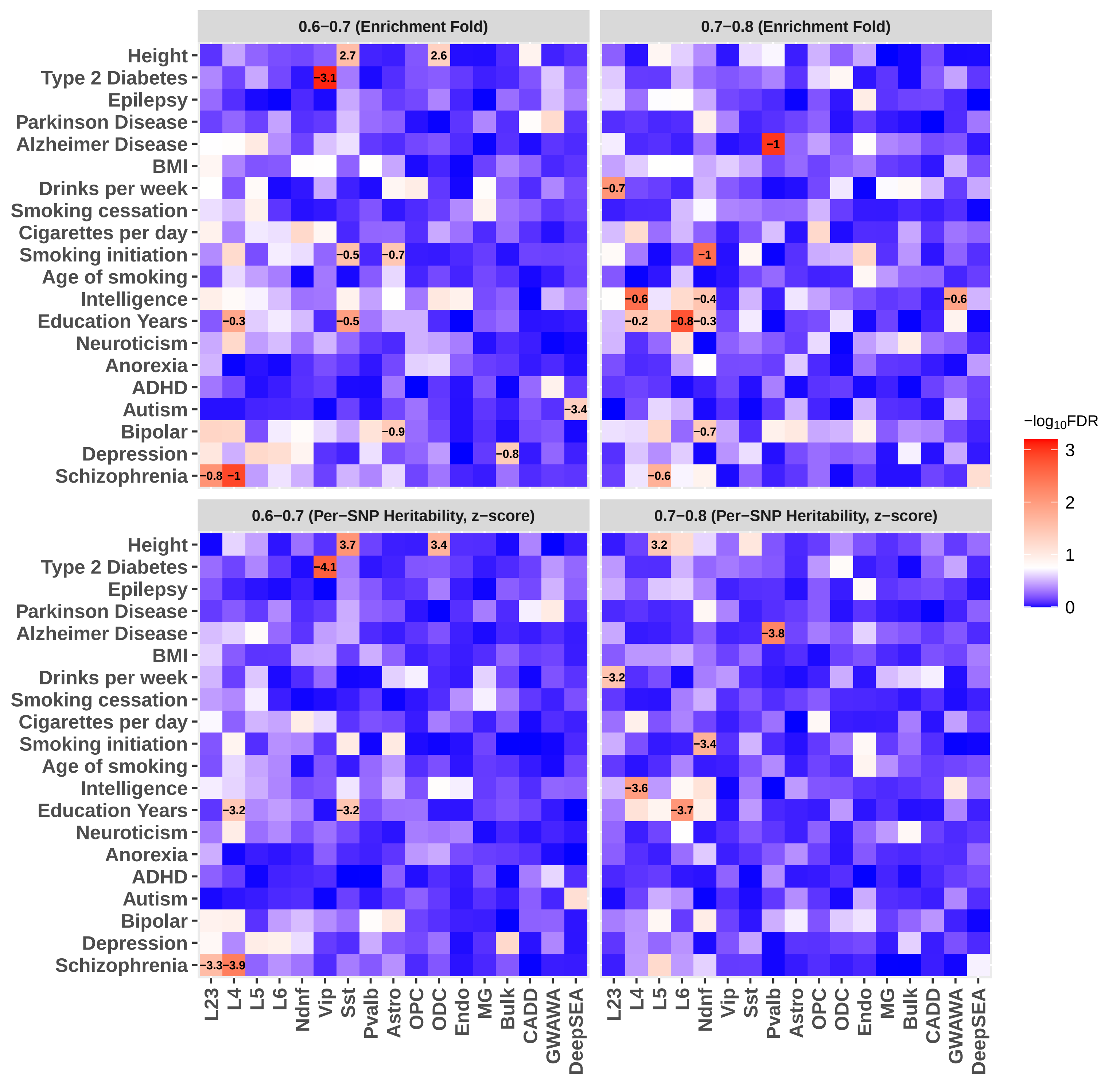

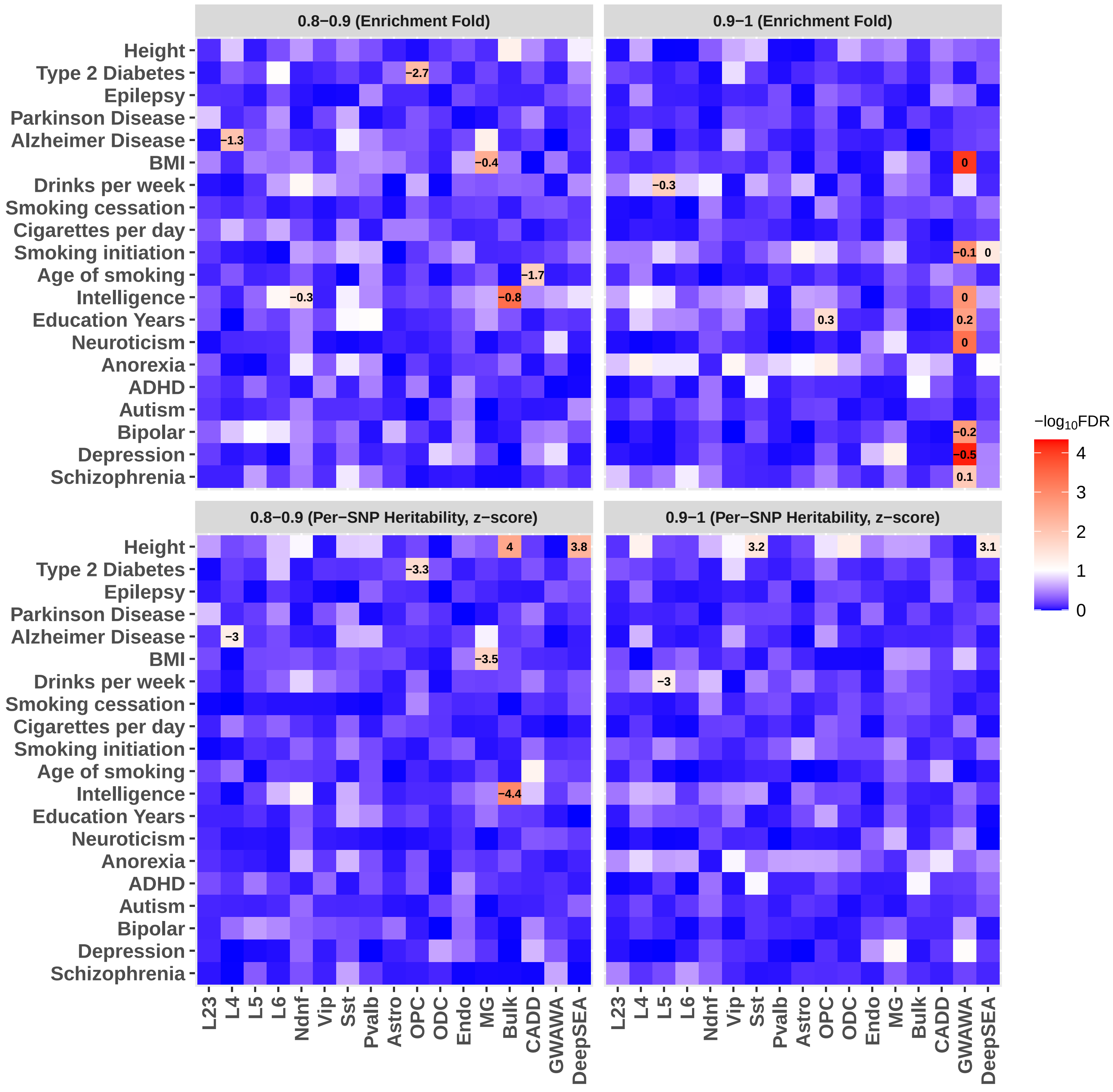
